# Cultural Innovations Influence Patterns of Genetic Diversity in Northwestern Amazonia

**DOI:** 10.1101/347336

**Authors:** Leonardo Arias, Roland Schröder, Alexander Hübner, Guillermo Barreto, Mark Stoneking, Brigitte Pakendorf

## Abstract

Human populations often exhibit contrasting patterns of genetic diversity in the mtDNA and the non-recombining portion of the Y-chromosome (NRY), which reflect sex-specific cultural behaviors and population histories. Here, we sequenced 2.3 Mb of the NRY from 284 individuals representing more than 30 Native-American groups from Northwestern Amazonia (NWA) and compared these data to previously generated mtDNA genomes from the same groups, to investigate the impact of cultural practices on genetic diversity and gain new insights about NWA population history. Relevant cultural practices in NWA include postmarital residential rules and linguistic-exogamy, a marital practice in which men are required to marry women speaking a different language.

We identified 2,969 SNPs in the NRY sequences; only 925 SNPs were previously described. The NRY and mtDNA data showed that males and females experienced different demographic histories: the female effective population size has been larger than that of males through time, and both markers show an increase in lineage diversification beginning ~5,000 years ago, with a male-specific expansion occurring ~3,500 years ago. These dates are too recent to be associated with agriculture, therefore we propose that they reflect technological innovations and the expansion of regional trade networks documented in the archaeological evidence. Furthermore, our study provides evidence of the impact of postmarital residence rules and linguistic exogamy on genetic diversity patterns. Finally, we highlight the importance of analyzing high-resolution mtDNA and NRY sequences to reconstruct demographic history, since this can differ considerably between males and females.

## INTRODUCTION

Uniparentally inherited mitochondrial DNA (mtDNA) and the non-recombining portion of the Y-chromosome (NRY) have been used extensively to study human population history. Their sex-specific mode of inheritance is useful for contrasting the maternal vs. paternal history of populations. Previous studies have found that human populations generally exhibit larger genetic differences for the NRY than for mtDNA (Kayser et al. 2003; Lippold et al. 2014; Seielstad et al. 1998). Several explanations revolving around cultural practices have been proposed to account for these differences. First, women migrate more often than men: around 70% of human societies are characterized by patrilocality (Burton et al. 1996), meaning that men generally remain in their birthplace, leading to an increase of the genetic differences in the NRY among populations. In contrast, the movement of women among populations results in a reduction of the genetic differences in the mtDNA (Gunnarsdottir et al. 2011; Hamilton et al. 2005; Heyer et al. 2012; Jobling and Tyler-Smith 2003; Marchi et al. 2017; Oota et al. 2001; Seielstad et al. 1998; Verdu et al. 2013). Second, some studies have claimed that these differences reflect a disparity between male and female effective population sizes (Ne), with the female Ne being larger than that of males. This might be a consequence of differential male reproductive success (i.e., many fewer males than females having offspring) (Heyer et al. 2012; Tang et al. 2002; Wilder et al. 2004). In addition, subsistence strategies have been shown to influence levels of genetic diversity within populations, with agriculturalist groups exhibiting higher genetic diversity than hunter-gatherer groups. This is probably a consequence of population expansions experienced by the former and contractions undergone by the latter (Aime et al. 2013; Excoffier and Schneider 1999; Patin et al. 2014).

Northwestern Amazonia (NWA) is particularly interesting in this regard, since humans have successfully adapted to its remarkable array of ecosystems since at least 10,000 years ago (Aceituno et al. 2013; Piperno 2011), and human societies inhabiting this region exhibit great diversity in terms of languages, subsistence strategies, marital practices, and residential patterns. Although rich ethnographic descriptions exist for the region that highlight the diversity of Amazonian societies (Chernela 2010; Hugh-Jones 1979; Koch-Grünberg 1995; Sorensen 1967; Steward 1949), there is a general lack of studies that address the impact of these cultural practices on the patterns of genetic variation, as well as studies that reconcile the patterns of variation at a local scale to the patterns of genetic variation observed for the Americas in general.

In order to address these issues, we have investigated the genetic diversity present in a comprehensive sample covering the extant ethnolinguistic diversity of NWA, including populations from different language families that have different subsistence strategies and different marital and other cultural practices. For example, some of the groups included in our study engage in a marital practice known as linguistic exogamy (Sorensen 1967; Stenzel 2005), in which men are required to marry women speaking a different language. We have employed next generation DNA sequencing methods to assess the levels of diversity in the mtDNA and NRY, thereby avoiding the methodological differences that have characterized many previous studies. With these data, we aim to investigate how the patterns of variation in Native American populations from NWA are affected by different cultural practices. In addition, we investigate how the sex-specific demographic patterns inferred from these two markers contribute to a reconstruction of the human population history of NWA.

## RESULTS

We generated 284 NRY sequences from Native American individuals from NWA, using a hybridization capture method that covers a region of ~2.3 Mb. The average coverage per sample was 37.37 +/− 15.55, which was reflected in the relatively low number of missing sites (mean = 3.99) (Supplementary Table 1). We identified 2,969 SNPs in the sequences analyzed, of which only 925 have been described in the International Society of Genetic Genealogy (ISOGG) database (www.isogg.org accessed on 22.02.2018) and assigned to particular lineages. Thus a total of 2,044 previously uncharacterized SNPs are reported here that allow a higher level of discrimination among sequences.

We compared the patterns of genetic variation between the NRY and mtDNA in 17 NWA ethnolinguistic groups, using a subset of the mtDNA genomes previously reported in Arias et al (2018). All mtDNA lineages belonged to autochthonous Native American haplogroups, namely A2, B2, C1, or D1. In contrast, the NRY lineage diversity was dominated by haplogroup Q1, the main haplogroup observed among Native Americans, which in our dataset reached a frequency of 91 % (Table 1). Within Q1 we identified a Cocama individual who exhibited a divergent Q1 sequence that marks the first split of the tree (Supplementary Figure 1) at 27.8 kya [95% highest posterior density (HPD) = 20.6-34.3 kya]. The sequence was classified as haplogroup Q1b1a1 since it carries derived alleles at the defining SNPs L275 and L612. This lineage, which has not been previously reported in Native Americans, occurs at high frequency in Central, West, and South Asia (Balanovsky et al. 2017), and the estimates of the divergence between Q1a and Q1b are around 30-32 kya (Kivisild 2017; Poznik et al. 2016). However, based on the sequence of just one individual it is difficult to determine if this lineage can be considered autochthonous in Native Americans, or if instead it reflects post-contact admixture. Additionally, haplogroup C2 (previously called C3), the other confirmed autochthonous lineage in the Americas (Roewer et al. 2013), was observed at very low frequency: only three individuals (1%) exhibited this haplogroup, two from the Guayabero and one from the Mur-Uitoto group. Furthermore, 18 individuals (8%) from the groups included in the population-based analyses carried lineages that are commonly regarded as the product of recent European or African admixture. Of these, haplogroup R1 was the most frequent, accounting for five percent of the total sequences. These observations are in agreement with previous studies that indicate that the patterns of admixture in American populations have been sex-biased (Mesa et al. 2000; Sans 2000), particularly during colonial times, involving primarily European men and indigenous women.

**Table 1.**
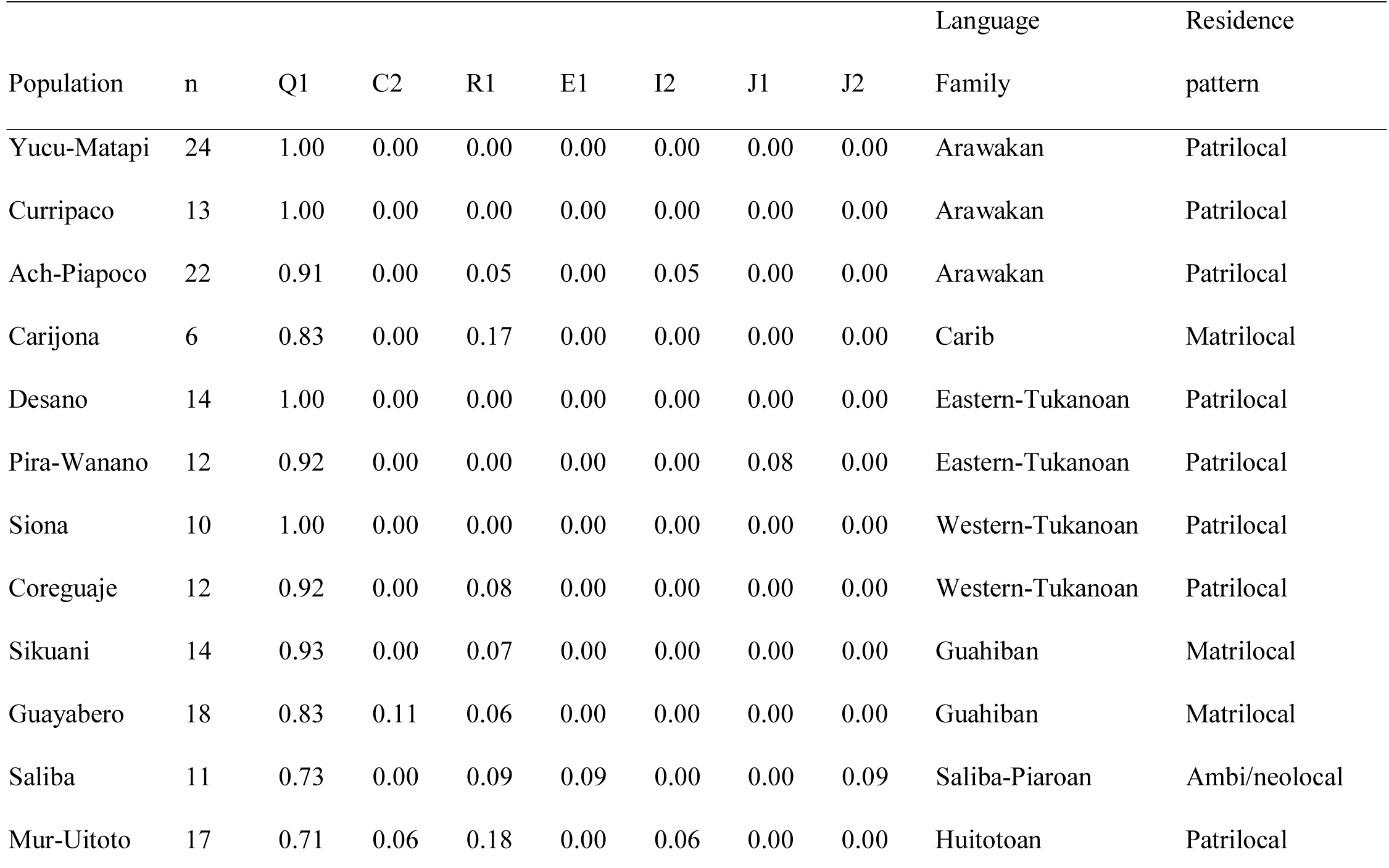
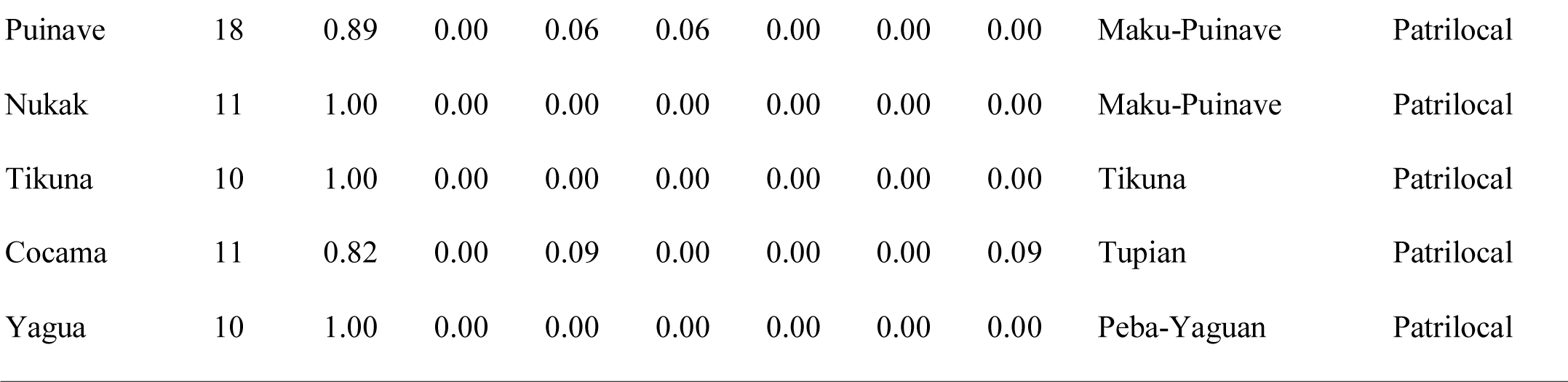
NRY haplogroup frequencies for the 17 NWA groups included in the population-based analyses

### Molecular diversity

The patterns of NRY and mtDNA genetic diversity varied among NWA populations. The Eastern Tukanoan Desano and Pira-Wanano stand out in having much lower than average Y-chromosomal gene diversity in conjunction with higher than average values of mtDNA gene diversity (Figure 1A). In contrast, the Siona and the Ach-Piapoco had higher than average NRY diversity and lower than average mtDNA diversity, while the Mur-Uitoto, Tikuna, and Cocama had higher than average gene diversity values for both markers; the hunter-gatherer groups Nukak and Sikuani had lower than average gene diversity values for both markers. In addition to low gene diversity values for the NRY, the Eastern Tukanoan groups also exhibited much lower than average values of the mean number of pairwise differences (MPD) for the NRY (6.6 in the Desano and 3.2 in the Pira-Wanano as opposed to 37.9 differences on average), whereas their mtDNA MPD values were in the average range. In contrast, the Guayabero and Mur-Uitototo showed the highest values of NRY MPD. However, when the C2 sequences were excluded, their values were within the range of the other populations (Figure 1B, Supplementary Table 2).

**Figure 1.**
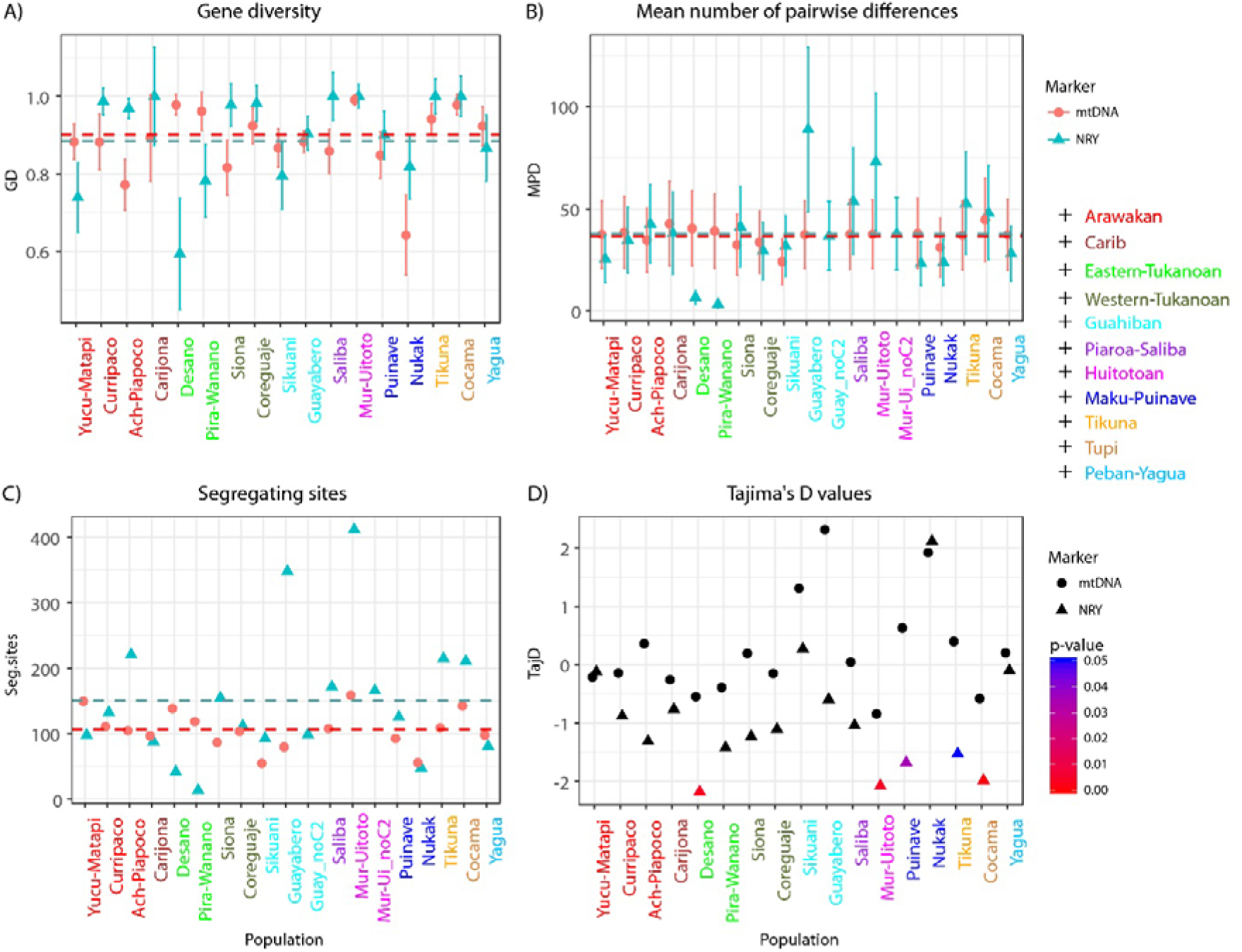
Molecular diversity indices for the mtDNA and NRY in 17 groups of NWA. A) Gene diversity. B) Mean number of pairwise differences. C) Number of segregating sites. D) Tajima’s D values. Dotted lines represent the average diversity values across all groups. Population names are color-coded by language family as indicated in the legend. Values for the NRY after removing sequences belonging to haplogroup C2 are shown next to Guayabero and Mur-Uitoto, the only groups carrying these sequences.

The number of segregating sites varied considerably among groups for both mtDNA (range from 55 to 159) and for the NRY (range from 14 to 412). In the NRY we observed five populations (Ach-Piapoco, Guayabero, Mur-Uitoto, Tikuna, and Cocama) with more than 200 segregating sites and three populations (Desano, Pira-Wanano, and Nukak) with less than 50 segregating sites. The high values in Guayabero and Mur-Uitoto are again explained by the presence of sequences belonging to haplogroup C2; when we excluded these sequences, the number of segregating sites dropped to 99 in Guayabero, although it remained high for Mur-Uitoto (166 segregating sites; Figure 1C). At the same time, the high numbers of segregating sites in the Ach-Piapoco, Tikuna, and Cocama might be an indication that these groups have a complex population history with heterogeneous origins.

Tajima’s test of selective neutrality (Figure 1D) showed more negative D values for the NRY than for mtDNA, and these were significantly negative (P-value < 0.05) in five populations, namely Desano, Mur-Uitoto, Puinave, Tikuna, and Cocama. In contrast, none of the D values estimated from the mtDNA were significant, not even the large positive values found for the Guayabero (2.3), Nukak (1.9), and Sikuani (1.3). Negative Tajima’s D values are indicative of population expansion, while positive values suggest population contractions. As the Nukak and Sikuani are the only groups exhibiting positive Tajima’s D values for both the mtDNA and NRY, they may have undergone a population contraction. In contrast, the lower D values observed in the NRY than in the mtDNA suggest that most of the NWA populations have undergone recent population expansions in the paternal line.

The comparisons of the patterns of genetic variation between matrilocal and patrilocal populations are shown in Figure 2. We obtained the distribution of the pairwise Φ_ST_s, the gene diversity, and the MPD by sampling all possible combinations of triplets of patrilocal populations and estimating, for every triplet, the mean value of each statistic. After correcting for multiple comparisons, there were no significant differences between patrilocal and matrilocal groups in any of the values. However, the distributions of the statistics were compatible with the expectations for patrilocality vs. matrilocality (Figure 2): only 2% of the patrilocal triplets exhibited larger MPD values than the matrilocal groups for the NRY, while for mtDNA 90% of patrilocal triplets exhibited larger MPD values than matrilocal groups. Moreover, 74% of the patrilocal triplets exhibited smaller pairwise Φ_ST_ values for the mtDNA than matrilocal groups, whereas 95% of patrilocal groups exhibited larger Φ_ST_ values for the NRY than matrilocal groups.

**Figure 2.**
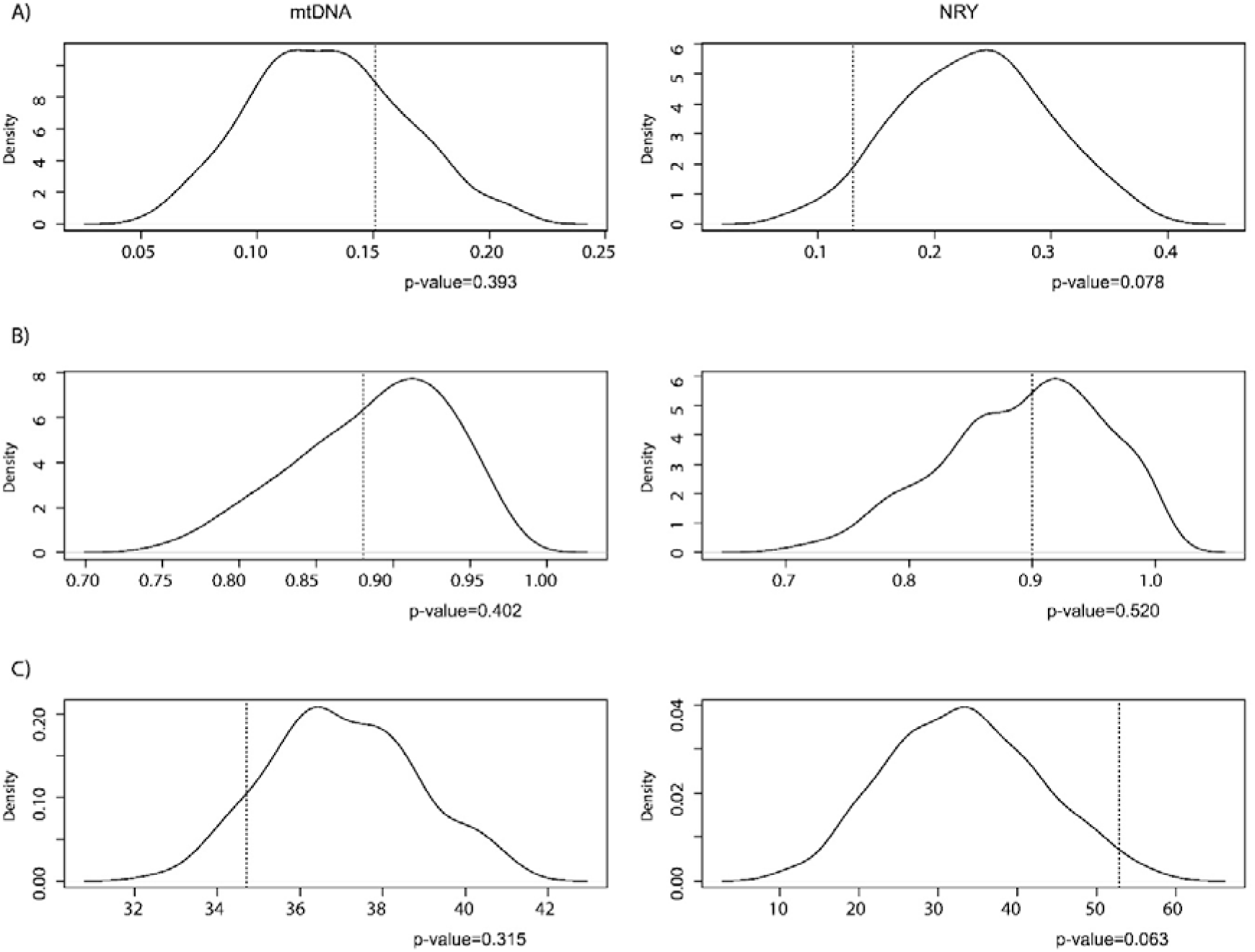
Distribution of the mean values for: A) Φ_ST_s, B) gene diversity, and C) MPD for all possible triplets (n=286) of patrilocal groups for the mtDNA (left side) and NRY (rightside). The mean values for matrilocal groups are shown as dotted lines. One-side p-values reported after Benjamin-Hochberg correction for multiple comparisons.

### Genetic structure and genetic distances

The AMOVA results (Table 2) showed that the percentage of between-population differences was higher for the NRY (27.24 %) than for the mtDNA (12.79 %), indicating that there is more genetic structure for the NRY and therefore larger genetic distances among populations, as we show below. In addition, the AMOVA results were consistent with our previous findings (Arias et al. 2018) that indicate that the distribution along rivers is a better predictor of the genetic structure in NWA than the general geographical location or the linguistic affiliation of groups (Table 2). This is particularly true for the NRY, since the component of variation due to differences among groups defined by settlement along rivers was highly significant and larger than the component of variation due to differences between populations within groups.

**Table 2.** AMOVA results

|  | Among groups | Within groups | Within populations | # groups |
| --- | --- | --- | --- | --- |
| **mtDNA** |  |  |  |  |
| Language | -3.79 | 16.37** | 87.42** | 11 |
| Geography | -0.83 | 13.53** | 87.30** | 6 |
| Rivers | 4.15* | 8.83** | 87.02** | 10 |
| Global $\Phi_{ST}$ | 12.79 | | | |
| **Y-chromosome** |  |  |  |  |
| Language | 5.77 | 21.81** | 72.42** | 11 |
| Geography | 6.71* | 21.23** | 72.06** | 6 |
| Rivers | 14.45** | 13.35** | 72.2** | 10 |
| Global $\Phi_{ST}$ | 27.24 | | | |

We used the pairwise Φ_ST_ values as a measure of the genetic distances among populations. Pairwise distances were generally lower for mtDNA and fewer values were significantly different from zero (p-value < 0.05) than for the NRY (Supplementary Figure 2). A visual representation of the matrices of pairwise Φ_ST_ distances are presented in the MDS plots (Supplementary Figure 3), where we observed different genetic relationships among populations for the mtDNA and the NRY. For instance, the Eastern Tukanoan groups Desano and Pira-Wanano were located in the center of the plot, together with several other groups, for the mtDNA, but were clearly differentiated based on the NRY (Supplementary Figure 3), and there was no obvious tendency for groups from the same language family to cluster together. Finally, a Mantel test revealed no significant correlations between genetic distances based on mtDNA and NRY sequences, nor between genetic and geographic distances (Supplementary Figure 4).

The shared haplotypes between pairs of populations provide information about their relationships, which can be due to common ancestry and/or gene flow between populations. Even though there was less sharing of haplotypes both between and within populations for the NRY in comparison to the mtDNA (Figure 3) — which can be explained by the differences in the amount of sequence and the mutation rate between the NRY and the mtDNA—the fact that there are shared NRY haplotypes between populations is noteworthy. According to the mutation rate used here of 7.6 × 10^−10^ subs/bp/year (Fu et al. 2014), for the sequenced region of ~2.3 Mb we expect to observe, on average, one substitution every 572 years, indicating that shared NRY haplotypes reflect recent contact or very recent population divergences; they thus further confirmed the contact events that we described previously based on mtDNA alone (Arias et al. 2018). For instance, the Guayaberos, Sikuani, Nukak, Puinave, Saliba, Ach-Piapoco, and Curripaco, who inhabit several tributaries of the Orinoco River, shared both identical and closely related mtDNA and NRY haplotypes (Figure 3 and Supplementary Figure 5), indicating that these groups have interacted for some time (Arias et al. 2018). This is additionally supported by ethnohistoric accounts of the existence of contact among the groups living along the Orinoco River and its tributaries (Morey and Morey 1980; Rey Fajardo 1974).

**Figure 3.**
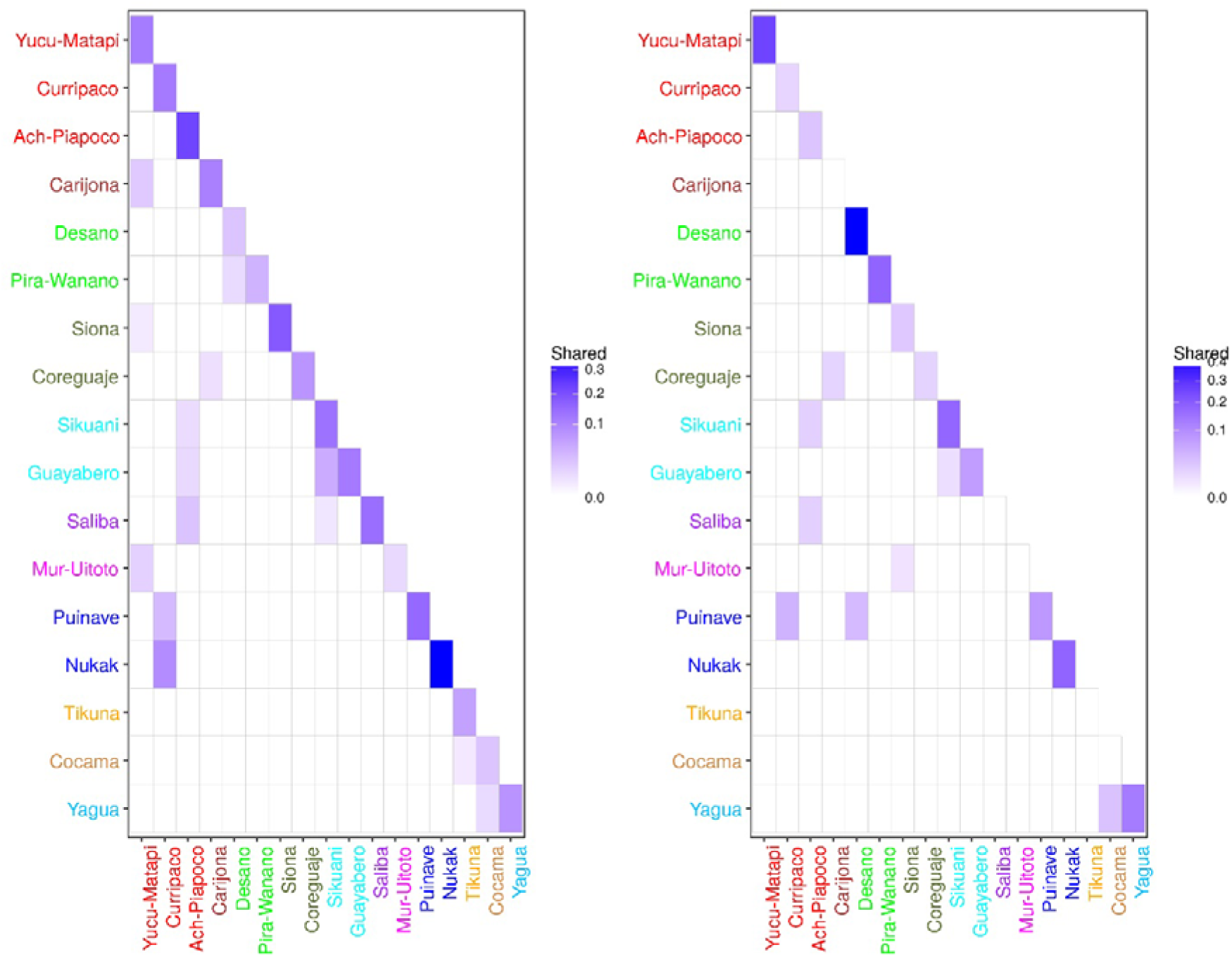
Shared haplotypes between pairs of populations for the mtDNA (left) and NRY (right).

### Phylogenetic and demographic reconstructions

The estimation of coalescence events through time and the phylogenies for the NRY haplogroup Q1 sequences and the mtDNA are depicted in Figure 4 and Supplementary Figures 1 and 6, respectively. The first split of the Q1 phylogeny separated a single individual belonging to subhaplogroup Q1b1a1 from the bulk of sequences belonging to haplogroup Q1a, which showed an initial event of diversification ~13-14 thousand years ago (kya). This diversification is evident in the number of coalescence events that happen around that time (Figure 4). An additional accumulation of coalescence events was observed after 5 kya, with 50% of all coalescence events happening during the last ~1 kya (Figure 4). The mtDNA also showed two regions of high accumulation of coalescence events (Figure 4), similar to what was found for the NRY. One such accumulation starts ~17.5 kya and the second one starts ~5 kya, with 50% of all coalescence events taking place in the last ~1 kya.

**Figure 4.**
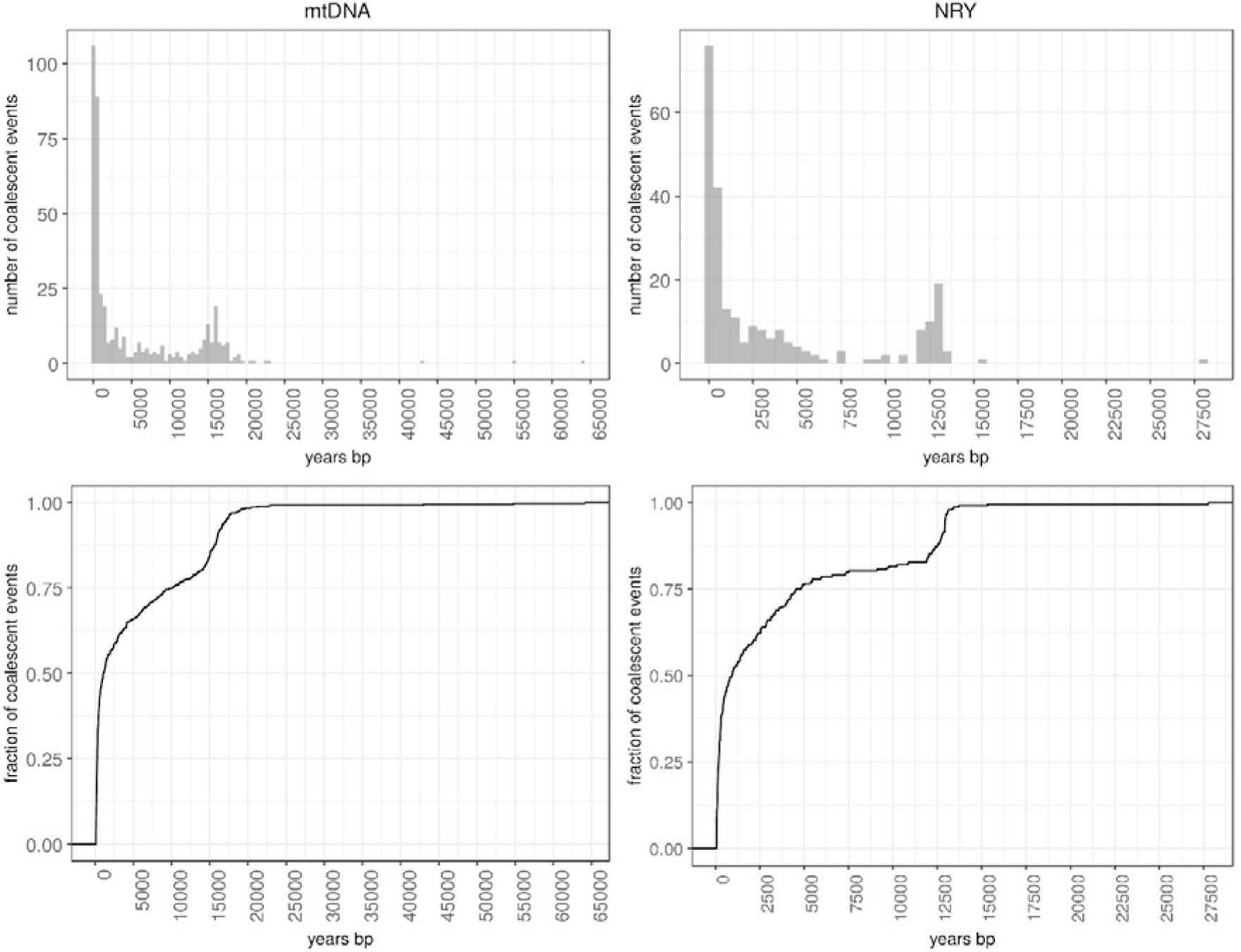
Histograms and cumulative fraction of the number of coalescent events through time for the mtDNA (left) and NRY (right).

The BSP demographic reconstructions for the mtDNA and the NRY are shown in Figure 5. Both markers exhibited striking increases in the effective population size (Ne). However, there were differences in the estimates of Ne, the magnitude of increase, and the dates for the start of the population expansion. The mtDNA exhibits a single signal of population expansion starting ~17 kya, while in the NRY we observed an initial expansion that starts at ~13.5 kya and a more recent expansion of smaller magnitude at ~3.5 kya, preceded by a reduction in Ne after 10 kya. Furthermore, the mtDNA exhibits larger Ne through time, being double the size of the NRY at the peak of both markers. The older signal of population expansion in the NRY was also observed in the network of haplotypes belonging to haplogroup Q1, which showed a star-like shape suggestive of a rapid diversification of lineages (Supplementary Figure 5). We estimated the date for this diversification around 13.6 +/- 0.6 kya, based on the rho statistic as implemented in the software Network 4.6.1.5 (http://www.fluxus-engineering.com), using the substitution rate reported by Fu et al. 2014 (7.6 × 10^−10^subs/bp/year).

**Figure 5.**
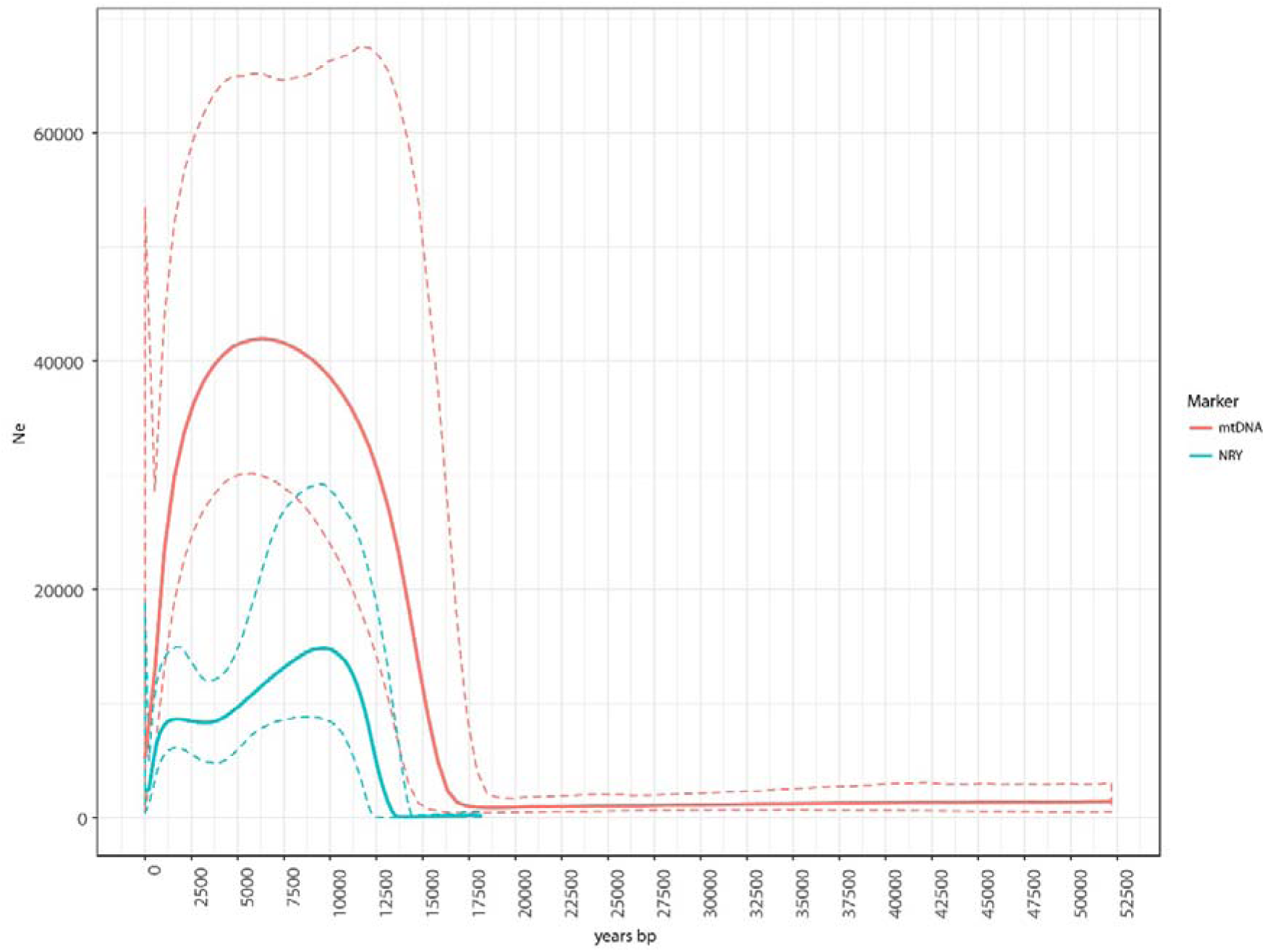
BSPs for the mtDNA and NRY sequences from NWA. The dotted lines indicate the 95% HPD intervals. Ne was corrected for generation time according to (Fenner 2005), using 26 years for mtDNA and 31 years for NRY.

We further investigated the differences between the BSPs for both markers. Although both the mtDNA and the NRY exhibit similar signals of recent coalescence events (Figure 4), the BSPs differ in that the NRY, but not the mtDNA, shows a recent expansion. A possible explanation is violation of the panmixia assumption, as it has been shown that coalescence-based methods, including BSP, can then produce misleading results (Grant 2015; Heller et al. 2013). That is, locally sampled populations that are interconnected by low levels of migration (a structured population) typically exhibit genealogies that resemble those of panmictic populations that have declined in size (Heller et al. 2013). One way to address this issue is to use different sampling strategies (see Methods). The BSP results of the “pooled” and “scattered” strategies consistently identified the old signal of population expansion in both mtDNA and NRY, while some local populations showed a flatter trajectory of the Ne through time (Supplementary Figures 7 and 8). Moreover, we observed that the BSPs from the pooled sampling strategy of the NRY, but not of the mtDNA, consistently showed the signal of a recent increase in population size between 2.5-3.5 kya (Supplementary Figure 7). This suggests that this increase was not due to violation of the panmixia assumption, but rather indicates a true male-biased expansion.

### Patterns of genetic diversity at the local vs. continental scale in Native Americans

Our comparisons of the patterns of genetic diversity at different geographic scales are shown in Figure 6. We observed differences in the levels of within population diversity (i.e., nucleotide diversity) and between population differentiation (i.e., Φ_ST_s) between the mtDNA and the NRY. First, the nucleotide diversity in the mtDNA was larger than in the NRY, without significant differences between the local and continental levels. Second, our estimates of the amount of between-population differences in the Americas contrast with the findings of Lippold et al. (2014), who reported much larger between-population differences for the mtDNA (Φ_ST_ = ~ 0.7) than for the NRY (Φ_ST_ = ~ 0.2) in a sample of 22 individuals from the CEPH Human Genome Diversity Panel. In contrast, we observed in our continental dataset (composed of 77 individuals from four countries, see Materials and Methods) a much smaller degree of differentiation for the mtDNA (Φ_ST_ = 0.08 averaged over five replicates) than that found by Lippold et al., while the NRY Φ_ST_ value (0.17) was comparable (Supplementary Table 3). Furthermore, we observed differences in the mtDNA between the Americas (Φ_ST_ = 0.08) and NWA (Φ_ST_ = 0.13) and much larger differences for the NRY between NWA groups (ST = 0.27) and groups sampled at a continental scale (Φ_ST_ = 0.17).

**Figure 6.**
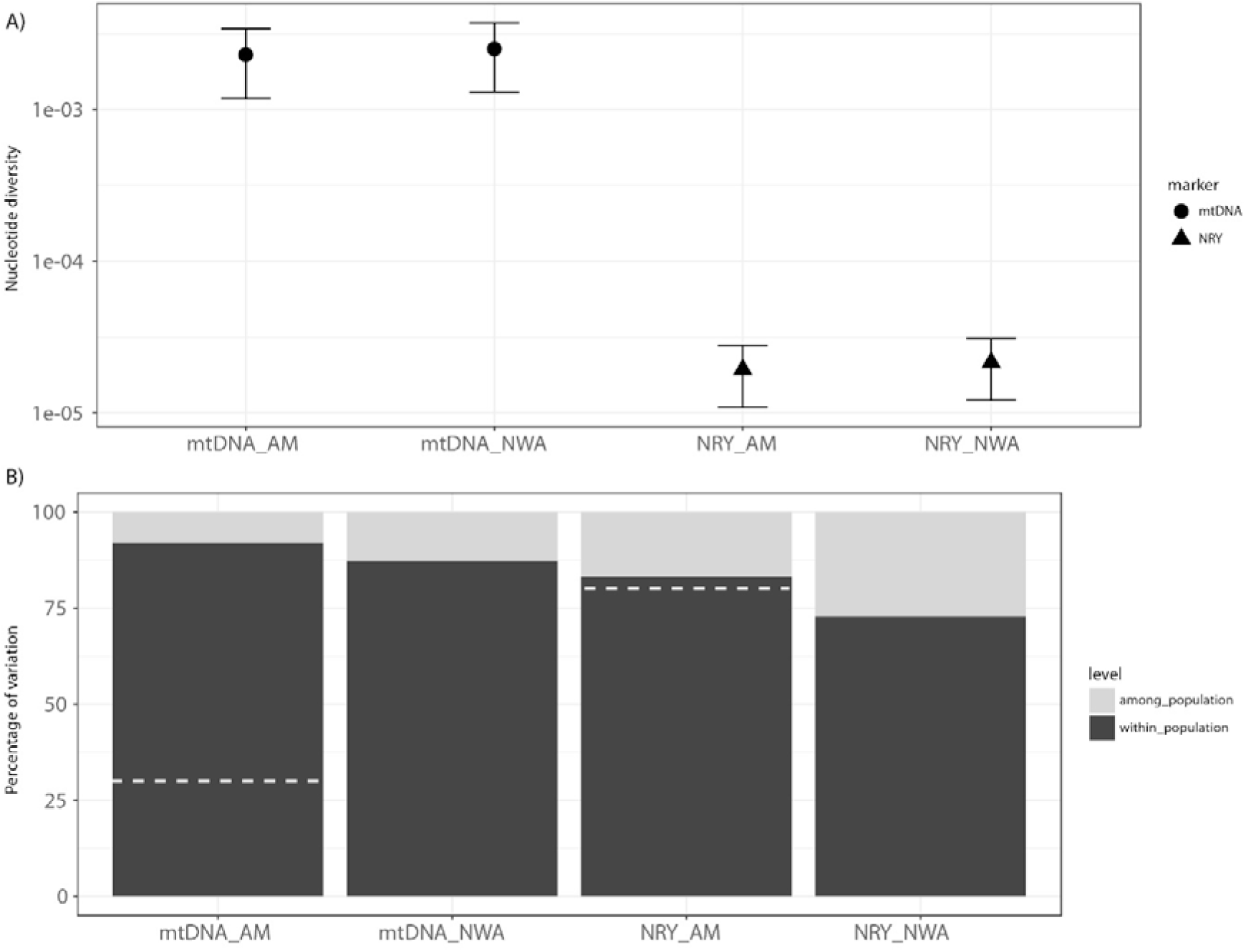
A) Nucleotide diversity and B) AMOVA results in NWA and a sample of the Americas (AM) for the mtDNA and the NRY. Dotted lines represent the percentage of among population variation reported by Lippold et al. (2014) for the Americas.

## DISCUSSION

We have investigated the patterns of genetic variation in a region of ~2.3 Mb of the Y-chromosome and compared them to complete mitochondrial genomes in a large sample of ethnolinguistic groups from NWA. This approach overcomes the drawbacks from previous studies regarding the quality and quantity of the data from which the levels of diversity in the mtDNA (typically sequences of the hypervariable segments of the control region) and in the Y-chromosome (typically biallelic SNPs and/or Y-STRs) have been assessed. We have identified 2,044 previously uncharacterized NRY SNPs, which provide new insights about NRY genetic diversity in Native Americans.

### The impact of cultural practices on patterns of genetic diversity across groups

Human populations in NWA exhibit differences in terms of cultural practices. In this study we have investigated how the patterns of genetic variation are impacted by cultural practices, in particular postmarital residence patterns and linguistic exogamy; these are discussed in the following sections.

### Postmarital residence patterns

Patrilocal groups are expected to exhibit lower levels of within-population genetic diversity and higher levels of between-population divergence (i.e. high pairwise Φ_ST_s) for the NRY, and higher levels of within-population diversity and lower levels of divergence for the mtDNA; matrilocal groups are expected to show the opposite pattern (Oota et al. 2001). However, as found previously in other regions of the world (Ascunce et al. 2008; Gunnarsdottir et al. 2011; Kumar et al. 2006), our data do not conform to the patterns expected for these postmarital residence practices. Only three populations out of 13 that are classified as patrilocal, namely, the two Eastern Tukanoan groups Desano and Pira-Wanano and the Arawakan group Yucu-Matapi, showed lower values of NRY gene diversity than two of the matrilocal groups, namely the Carijona and Guayabero, while only the Desano and Pira-Wanano exhibited larger mtDNA gene diversity values. The Sikuani, also classified as matrilocal, showed lower NRY gene diversity values, in contrast to what would be expected from their residence pattern. In general, there is considerable heterogeneity among patrilocal groups in the gene diversity values (Figure 1A). For instance, the patrilocal groups Ach-Piapoco and Siona showed a pattern that looks more like matrilocality, namely high NRY and low mtDNA gene diversity. Similarly, the three groups classified as matrilocal—Carijona, Sikuani, and Guayabero—did not significantly differ from the gene diversity values observed for many of the patrilocal groups (Figure 1A).

Since patrilocal groups in our study are more heterogeneous and considerably outnumber the matrilocal groups, the resampling approach allowed for a more reliable comparison between residence patterns. This analysis detected that patrilocal groups have lower MPD values for the NRY than matrilocal groups (p-value = 0.063). In the other comparisons, we also observed a tendency in the distribution of the evaluated statistics that is compatible with the general expectations for patrilocality vs. matrilocality (Figure 2). Thus, patrilocal groups showed on average smaller pairwise Φ_ST_ and larger MPD for the mtDNA than matrilocal groups. Meanwhile, we observed the opposite pattern for the NRY, that is, larger pairwise Φ_ST_s and smaller MPD in the patrilocal than in the matrilocal groups. In contrast, we did not find significant differences in the gene diversity values between matrilocal and patrilocal groups (Figure 2). In summary, we can say that NWA populations show considerable heterogeneity in the levels of genetic variation, which cannot be fully explained by a simple distinction between matrilocal and patrilocal groups. Rather, this is just one of the factors at play in structuring the observed genetic variation. Alternatively, our results might be an indication that in practice there is more flexibility in postmarital residence rules than what is considered the cultural norm in theory (Hamilton et al. 2005; Walker 2015).

### Linguistic exogamy

One of the most striking patterns among NWA populations is observed in the two Eastern Tukanoan groups, who show higher than average mtDNA diversity and lower than average NRY diversity (Figure 1A), as well as an extreme reduction of the MPD for the NRY (Figure 1B). In our previous study (Arias et al. 2018), which included more Eastern Tukanoan groups, we found that five out of the six groups showed higher than average mtDNA diversity. Eastern Tukanoan groups represent a prime example of the impact of cultural practices on the patterns of genetic diversity. Specifically, Eastern Tukanoans practice linguistic exogamy, a cultural norm in which marriages are required to occur between individuals speaking different languages. This system promotes the migration of women among ethnolinguistic groups, since it is reinforced by patrilocality and patrilineality, in which married men continue to live in their fathers’ territory and the individual’s identity is determined by the father’s ethnolinguistic identity (Sorensen 1967; Stenzel 2005). For instance, Stenzel (2005) reports that in the Vaupes region 75-90% of the married men continue to reside in the sub-region in which they lived before marriage, while this is true for only 50-58% of the women in the same region. This situation gives rise to the high levels of mtDNA and low levels of NRY genetic diversity found in this study for the Eastern Tukanoan Desano and Pira-Wanano.

### Patterns of genetic diversity at the local vs. continental scale in Native Americans

Previous studies have shown that human populations generally exhibit larger genetic differences among populations in the NRY than in the mtDNA (Fagundes et al. 2002; Kayser et al. 2003; Marchi et al. 2017; Nasidze et al. 2004; Oota et al. 2001; Pakendorf et al. 2007; Seielstad et al. 1998). However, Lippold et al. (2014) found differences in the patterns of genetic variation among continental regions, with the Americas, Oceania, and Africa exhibiting bigger between-population differences for mtDNA than for the NRY. We further explored this discrepancy by comparing a larger number of individuals from the Americas (n=77) in comparison to Lippold et al. (2014), who analyzed data for 22 individuals. In agreement with their results, we observed larger values of mtDNA nucleotide diversity than in the NRY. This could reflect differences in mutation rates between markers; however, previous studies (Poznik et al. 2016; Wilson Sayres et al. 2014) have shown that the differences in nucleotide diversity between the mtDNA and the NRY are maintained after correcting for differences in mutation rates.

In contrast to Lippold et al. (2014), our results showed smaller between-population differences for the mtDNA (Φ_ST_ = 0.08) than for the NRY (Φ_ST_ = 0.17) when comparing a larger number of sequences from the Americas (Figure 6). Thus, this region patterns as expected for human populations that are mostly patrilocal. Therefore, the much larger between-population differences for the mtDNA than for the NRY detected by Lippold et al. in the Americas are likely a result of the small sample size used in their study.

Furthermore, we found evidence that the patterns of genetic differentiation depend on the geographical scale of the study. The magnitude of between-population differentiation in the NRY compared to the mtDNA is smaller when looking at the continental scale than in NWA (Figure 6). This is in agreement with the findings of Wilkins and Marlowe (2006), who showed that the excess of between-population differentiation for the NRY in comparison to the mtDNA decreases when comparing more geographically distant populations. Heyer et al. (2012) and Wilkins and Marlowe (2006) have proposed that at a local scale the patterns of genetic diversity reflect cultural practices over a relatively small number of generations, whereas at a larger geographic scale the genetic diversity reflects old migration and/or old common ancestry patterns (Heyer et al. 2012; Wilkins and Marlowe 2006).

### The signals of an old and a recent diversification in NWA

Recent genetic evidence from contemporary populations and ancient DNA have provided evidence that the initial peopling of the Americas by modern humans occurred between 15-20 kya (Fagundes et al. 2008; Llamas et al. 2016; Moreno-Mayar et al. 2018; Raghavan et al. 2015). Once in the Americas, populations quickly expanded and spread, reaching the southern extreme of South America in a few thousand years. The archaeological site of Monte Verde in Chile, dated to ~14.5 kya, represents evidence of this rapid movement across the Americas (Dillehay et al. 2015). Our phylogenetic and BSP reconstructions are consistent with the signals of an old population expansion, happening between 13.5 kya (NRY) - 17 kya (mtDNA), in agreement with the proposed time frame for the peopling of the Americas.

The BSP plots and the diversity statistics indicate that overall the Ne of males has been smaller than that of females. One tentative explanation for this difference is that it reflects larger differences in reproductive success among males than among females. Some support for this explanation comes from the shape of the phylogenies (Supplementary Figures 1 and 6), since differences in reproductive success and the cultural transmission of fertility lead to imbalance phylogenies (Blum et al. 2006; Heyer et al. 2015). We estimated a common index of tree imbalance (Colless index) and calculated whether the mtDNA and NRY trees were more unbalanced than 1000 simulated trees generated under a Yule process (Bortolussi et al. 2006) (i.e. a simple pure birth process that assumes that the birth rate of new lineages is the same along the tree). We found that the NRY tree is more unbalanced than predicted by the Yule model (p-value=0.001), whereas the mtDNA tree is not significantly different from trees generated by the Yule model (p-value=0.628). It has been suggested that highly mobile hunter-gatherer societies, such as those typical of most of human prehistory, were polygynous bands (Dupanloup et al. 2003); similarly, nomadic horticulturalist Amazonian societies exhibit strong differences in reproductive success due to the common practice of polygyny, especially among community chiefs, whose offspring also enjoy a high fertility (Neel 1970; 1980; Neel and Weiss 1975).

Furthermore, a more recent expansion can be observed in the BSP based on the NRY, but not in the mtDNA BSP (Figure 5), indicating an expansion specifically in the paternal line. The reasons behind this recent male-biased population expansion, which starts ~3.5 kya, are as yet unclear. However, similar male-biased expansions have been observed in other studies using high-resolution NRY sequences (Batini et al. 2017; Karmin et al. 2015).

In addition, a recent event of lineage diversification is inferred from the accumulation of coalescent events for both mtDNA and the NRY (Figure 4). After ~ 5 kya there is evidence of a slight increase, and by ~1 kya a dramatic increase in the number of coalescent events, with around 50% of all coalescent events happening during the last 1,000 years. The expansion of widespread language families and/or archaeological cultures is often explained with the development of agriculture during the Neolithic (Bellwood et al. 2002; Diamond and Bellwood 2003; Gignoux et al. 2011; Renfrew 1999). However, the date for the recent diversification we detected in NWA is much later than the proposed dates for the domestication/management of plants in South America (~8 kya (Piperno 2011)). Therefore, we require a different explanation to account for the signal of diversification of mtDNA and NRY lineages (Figures 4) and the signal of population expansion detected in the NRY BSPs (Figure 5, Supplementary Figure 7). We propose that these results are likely to be the consequence of several interrelated factors: climate change, technological innovations, and the establishment of regional networks of exchange and trade.

Paleoenvironmental information indicates that during the transition from the Middle to Late Holocene (~4.2 kya) there was a major climatic change in tropical South America from drier conditions to increased rainfall (Cross et al. 2000; Marchant and Hooghiemstra 2004; Wanner et al. 2008). Sediment cores from lakes and pollen from terrestrial records in Amazonia indicate that this increase in rainfall promoted the spread of tropical rainforest over areas previously covered by savannas (Behling and Hooghiemstra 2000). We hypothesize that this major transition in the vegetation of Amazonia could have driven changes in subsistence strategies among human societies, as has been suggested by others (Iriarte et al. 2017). The archaeological record supports this transition, since it shows that large areas of tropical rainforest were modified by human activity, as indicated by changes in phytoliths assemblages and the presence of anthropogenic soils, a result of both increasing sedentism and agricultural activities (Clement et al. 2015; Levis et al. 2017). Especially by one kya, the archaeological record shows a sudden increase in the number of sites with human occupation with the presence of dark earths (Arroyo-Kalin 2010; Eden et al. 1984; Neves 2008). One crop that has been suggested as a candidate to allow this change in lifestyle is manioc (*Manihot esculenta*), especially the bitter variety, which grows well even in acidic and nutrient-poor soils, is more resistant to parasites, and produces larger and starchier tubers than the sweet manioc variety (Arroyo-Kalin 2010). However, bitter manioc tubers are highly toxic and require elaborate processing before consumption, including grating, washing, squeezing, and cooking. Therefore, the technological innovations that facilitated the utilization of bitter manioc roots, including the tools and utensils to process and cook the roots, supplied Amazonian societies with a constant and secure source of carbohydrates that resulted in increasing sedentism.

We additionally propose that women played an important role in the spreading of cultural innovations. Our data show a high level of migration of women among populations, as reflected in the low levels of population differentiation and the high amount of shared haplotypes in the mtDNA (Supplementary Figure 2 and 3; Figure 3). Manioc production is exclusively women’s work, and it includes planting, harvesting, processing and the preparation of different foods and drinks (Heckler 2004; Hugh-Jones 1979; Jackson 1983). Women are also important in the exchange of manioc landraces among groups, since in several ethnolinguistic groups of NWA a newly married woman receives several manioc varieties from her mother and grandmother as part of her dowry before she leaves for the community of her husband (Peña-Venegas et al. 2014). Any technological and horticultural innovations developed in one ethnolinguistic group would thus have been quickly spread to other groups by migrating women.

Furthermore, these cultural changes went hand in hand with the establishment of complex trade networks, where multilingualism was a common trait, and connected distant parts of Amazonia, the Andes and the Caribbean (Heckenberger and Neves 2009; Hornborg 2005; Vidal 1997; 2002). The last 4,000 years witnessed the geographic expansion of the major language families in different parts of South America, namely Arawakan, Tupi-Guarani, Carib, Ge, and Quechuan (Beresford-Jones and Heggarty 2011; Heckenberger 2002; 2013; Heggarty and Beresford-Jones 2012; Noelli 1998). Moreover, the analysis of the summed calibrated radiocarbon dates have shown the existence of a phase of population growth after ~5 kya (Goldberg et al. 2016). We have shown that this period is associated with increased diversification of both mtDNA and NRY lineages in NWA. Thus, what we observe in NWA may reflect processes that were happening in different parts of the continent, a cultural transition defined by Ford (1969) (Ford and Smithsonian 1969) as the “Formative Period”. The development of societies with increasing levels of complexity, that is, of societies living in permanent settlements and relying primarily on agriculture, producing sophisticated pottery and engaged in political, economic, and religious relationships with other societies, thus appears to have had a notable impact on the diversification of both mtDNA and NRY lineages in NWA.

Human prehistory is a complex phenomenon that can only be elucidated through the investigation of multiple lines of evidence from multiple disciplines. Our study has contributed to the understanding of the genetic history of the populations of NWA. By analyzing high-resolution mtDNA and NRY sequences, we have found that males and females have experienced different demographic histories. Cultural practices can account for some of the differences in the patterns of genetic diversity, for instance, by promoting differential reproductive success among individuals and determining the way the sexes disperse after marriage. We anticipate that the analysis of genome-wide data and ancient DNA studies of human remains will help fill in the remaining gaps in our knowledge.

## MATERIALS AND METHODS

### Sample collection

Saliva or blood samples were collected during several expeditions carried out by one of the authors (L.A.). Written informed consent was obtained from each participant and from the community leader and/or local/regional indigenous organizations after giving a full description of the aims of the study. All procedures were undertaken in accordance with the Declaration of Helsinki and the study was approved by the Institutional Review Committee on Human Ethics of the Universidad del Valle in Cali, Colombia, and the Ethics Commission of the University of Leipzig Medical Faculty. The total sample collection comprises 460 samples belonging to 40 different ethnolinguistic groups (see Arias et al. (2018) for details).

### DNA sequencing and sequence analysis

Double indexed DNA libraries (Kircher et al. 2012) were enriched for a region of 2.3 Mb of the non-recombining region of the Y-chromosome (NRY) via in-solution capture (Kutanan et al. 2018), following the SureSelect protocol from Agilent with modifications described in (Kircher et al. 2012). Three pools containing 90 samples each were prepared, and paired-end sequencing (read lengths 100 bp) was carried out on three lanes of the Illumina HiSeq 2500 platform; base-calling was performed with Bustard. Illumina adaptors were trimmed and reads starting from opposite directions were merged with leeHOM (Renaud et al. 2014a) to completely overlap paired sequences. Finally, sequences were de-multiplexed with the program deML (Renaud et al. 2014b) and aligned to the human reference genome *hg19* using BWA’s *aln* algorithm. All sequence pairs that aligned to the NRY regions defined by Poznik et al (2013) were retained (Poznik et al. 2013). Duplicate reads were removed using PicardTools *MarkDuplicates*, indel realignment was performed using GATK *IndelRealigner* (McKenna et al. 2010), and base quality was re-calibrated using GATK *BaseRecalibrator*. We identified single nucleotide variants (SNVs) using GATK *UnifiedGenotyper* v3.3-0 across all samples simultaneously, setting the parameter *ploidy* to 1 and using DBSNP build 138 as a prior position list. The identified SNVs were further filtered as previously described by Barbieri et al (2016) (Barbieri et al. 2016) and we obtained a set of 2969 SNVs. We imputed all samples with missing genotype information at any of these variant sites using BEAGLE (Browning and Browning 2013) and assigned Y chromosome haplogroups using yhaplo (Poznik et al. 2016).

The complete mtDNA genome sequences were taken from a previous study (Arias et al. 2018).

### Y-chromosome sequences

We generated NRY sequences from 284 individuals. From these we excluded seven “Mestizo” individuals (i.e., individuals with paternal origin from outside NWA) as well as 30 sequences belonging to non-autochthonous haplogroups. For the population-based analyses we excluded a further 32 sequences, since we restricted these analyses to populations with a sample size of at least 10 individuals for which we have both mtDNA and NRY sequence data. The sole exception was the Carijona (n=6), since it is the only Carib-speaking group living in NWA. Some ethnolinguistic groups were merged into single populations based on linguistic criteria when their population sizes were smaller than 10 individuals, as described previously (Arias et al. 2018). Thus, 215 NRY sequences and 330 complete mtDNA genome sequences from 17 ethnolinguistic groups were included in the population-based analyses. In contrast, for the phylogenetic and demographic reconstructions we included all the autochthonous sequences with the exception of three sequences belonging to haplogroup C2; these reconstructions were therefore based on 244 sequences belonging to haplogroup Q1. Seven populations included in Arias et al. (2018) were excluded due to insufficient numbers of NRY sequences. These are: the Eastern Tukanoan groups Tuka-Tatuyo, Siriano, Other-ET, and Tanimuka, as well as the three groups from the Andean foothills: Pasto, Kamentsa, and Inga. The population-based analyses included: the estimation of haplogroup frequencies (by counting); diversity values, AMOVA, and pairwise Φ_ST_ genetic distances, which were computed in Arlequin 3.5. (Excoffier and Lischer 2010); Multidimensional scaling (MDS) analyses based on pairwise genetic distance matrices, which were performed with the R package *MASS* (Venables and Ripley 2002); and the analysis of shared haplotypes, which was performed with in-house R scripts.

Furthermore, we investigated the impact of patrilocality and matrilocality on the patterns of genetic diversity. Our dataset contains 13 populations reported as patrilocal, three as matrilocal, and one as ambi/neolocal (Table 1). In order to control for the difference in the number of groups in each category and the heterogeneity among patrilocal groups, we devised a resampling strategy in which we sampled all possible triplets of patrilocal populations (286 combinations in total) and estimated for every sample the mean values of the following statistics: pairwise Φ_ST_ values, gene diversity and the MPD for the mtDNA and the NRY. We then compared these statistics to those observed among the matrilocal populations.

### Genetic structure and population relationships

For the AMOVA we defined clusters based on linguistic criteria, geographic proximity, and the distribution of populations along rivers; the latter is an important factor in structuring the mtDNA variation among NWA populations (Arias et al. 2018). For inferring the relationships among populations, we used pairwise Φ_ST_ values, as well as the proportion of shared haplotypes between populations. Additionally, we performed Mantel tests with the R package *ade4* (Dray and Dufour 2007) to evaluate whether there are significant correlations between pairwise genetic distances among populations for the NRY and the mtDNA, as well as between genetic distances and geographic distances. For the matrix of geographic distances we used the geographic coordinates of the locality that contained the majority of individuals for each ethnolinguistic group, and we calculated great circle distances between locations with the R package *geosphere* (Hijmans 2016).

### Phylogenetic inferences and demographic reconstructions

We reconstructed the phylogenies of the mtDNA and NRY sequences, using 428 complete mtDNA genome sequences reported by Arias et al (2018), and 244 NRY sequences belonging to haplogroup Q1. We estimated maximum clade credibility trees for both mtDNA and NRY sequences with BEAST 1.8.2 (Drummond et al. 2012). The best nucleotide substitution model was estimated with jModeltest 2.1.7 (Darriba et al. 2012), and we tested if the data best fit a strict clock or an uncorrelated log normal relaxed clock model using stepping-stone sampling (Baele et al. 2013) and Bayes factor analysis (Kass and Raftery 1995). Finally, we used the substitution rate of 7.6 × 10^−10^ per base pair per year reported by Fu et al (2014)(Fu et al. 2014) for the NRY, and substitution rates of 1.708 × 10^−8^ and 9.883 × 10^−8^ per base pair per year reported by Soares et al (2009) (Soares et al. 2009) for the coding and noncoding regions, respectively, of the mtDNA genome. In addition, a median-joining network of haplotypes for the NRY sequences was generated with the software Network 4.6.1.5 and visualized with Network Publisher 2.0.0.1 (http://www.fluxus-engineering.com).

We made inferences about changes in population size through time with Bayesian Skyline Plots (BSP) as implemented in BEAST 1.8.2 (Drummond et al. 2012). For this we used the same settings defined for the phylogenetic reconstructions (i.e., substitution model, substitution rate, clock model, etc.), but using the coalescent tree prior Bayesian skyline plot (BSP). We performed this analysis by haplogroup and by population. Furthermore, since violations of the panmixia assumption are known to lead to misleading results in coalescent-based methods (Grant 2015; Heller et al. 2013), we implemented the sampling strategies suggested by Städler et al. (2009), namely, “pooled” and “scattered”, and compared them with the results from the groups included in the population-based analyses, which corresponds to the “local” sampling scheme in (Städler et al. 2009). The “pooled” sample strategy consisted of randomly sampling four individuals from each population, while the “scattered” sample consisted of randomly sampling one individual from each population; each strategy was replicated ten times. This was performed for both the mtDNA and the NRY, and BSPs were obtained for each replicate.

### Patterns of genetic diversity at a local vs. continental scale in Native Americans

In order to investigate the effect of geographic scale on the patterns of genetic diversity, we compared several summary statistics at two levels: first, our local populations from NWA; and second, a comparative dataset of mtDNA and NRY sequences from the Americas, previously reported. The NRY comparative dataset was composed of 52 sequences from the Americas (Karmin et al. 2015; Mallick et al. 2016; Poznik et al. 2016), grouped by country of origin (i.e., Argentina n=12, Mexico n=15, and Peru n=25) and a random sample (n=25) of NRY sequences from NWA as representative of Colombia. We produced five NRY dataset replicates, by randomly sampling 25 NRY sequences from Colombia five independent times and merging with our 52 sequences from the literature. The mtDNA comparative dataset contained the same number of sequences by country as the NRY dataset (Argentina=12, Mexico=15, and Peru=25, Colombia=25). Five dataset replicates were generated by randomly sampling mtDNA sequences from a big collection of sequences from the literature (Achilli et al. 2013; Arias et al. 2018; Barbieri et al. 2017; Bodner et al. 2012; Cardoso et al. 2012; de Saint Pierre et al. 2012; Fagundes et al. 2008; Gomez-Carballa et al. 2012; Kumar et al. 2011; Lee and Merriwether 2015; Lippold et al. 2014; Mizuno et al. 2014; Perego et al. 2009; Perego et al. 2010; Tamm et al. 2007). We estimated the nucleotide diversity and performed an AMOVA considering two hierarchical levels, namely differences among populations and within populations. We compared the average values over the five replicates to the values observed in the NWA dataset.

### Data availability

Newly generated NRY sequences will be available upon acceptance of the manuscript.

## Acknowledgements

We greatly acknowledge the contribution of all sample donors, communities, community leaders, and regional indigenous organizations, without whose contribution this study would not be possible. We dedicate this article to Rafael Rodríguez, who assisted greatly with the fieldwork and passed away in September 2017. We also acknowledge Enrico Macholdt, Sandra Oliveira, Michael Dannemann, and Benjamin Peter for advice with data analysis. We thank Anna Paschall for her assistance with sample preparation, and Thiago Chacon, Zachary O’Hagan, and Jorge Rosés Labrada for their insights into the ethnohistory of Amazonia. B.P. acknowledges the LABEX ASLAN (ANR-10-LABX-0081) of Université de Lyon for its financial support within the program “Investissements d’Avenir” (ANR-11-IDEX-0007) of the French government operated by the National Research Agency (ANR). L.A. was supported by a graduate grant from COLCIENCIAS. This research was supported by funds from the Max Planck Society.

## Author contributions

B.P., M.S., and L.A. conceived and designed the project; L.A. collected samples in the field; R.S. assisted in data generation; A.H. assisted in merging NRY datasets from literature; G.B. contributed logistic support and reagents; L.A., M.S., and B.P. wrote the paper with input from all co-authors.

